# The transmembrane domain of DWORF activates SERCA directly; P15 and W22 residues are essential

**DOI:** 10.1101/2020.09.18.303644

**Authors:** Ang Li, Daniel R. Stroik, Samantha L. Yuen, Evan Kleinboehl, Razvan L. Cornea, David D. Thomas

## Abstract

The Ca-ATPase isoform 2a (SERCA2a) re-sequesters cytosolic Ca^2+^ into the sarcoplasmic reticulum (SR) of cardiac myocytes, enabling muscle relaxation during diastole. A central factor in heart failure is abnormally high cytosolic [Ca^2+^], resulting in pathophysiology and decreased cardiac performance. Therefore, augmentation of the SERCA2a Ca^2+^ transport activity is a promising therapeutic approach. A novel transmembrane peptide, dwarf open reading frame (DWORF), is proposed to enhance SR Ca^2+^ uptake and myocyte contractility by displacing the protein phospholamban (PLB) from its inhibitory site on SERCA2a. In the present study, we have developed several cell-based FRET biosensor systems for time-resolved FRET (TR-FRET) measurements of the protein-protein interactions and structural changes in SERCA2a complexes with PLB and/or DWORF. To test the hypothesis that DWORF competes with PLB to occupy the putative SERCA2a binding site, we transiently transfected DWORF into a stable cell line expressing SERCA2a labeled with green fluorescent protein (GFP, the FRET donor) and PLB labeled with red fluorescent protein (RFP, the FRET acceptor). We observed a significant decrease in FRET efficiency, consistent with a decrease in the fraction of SERCA2a bound to PLB. Functional analysis demonstrates that DWORF activates SERCA in both the presence and absence of PLB. Furthermore, using site-directed mutagenesis, we generated DWORF variants that do not activate SERCA, thus identifying residues that are necessary for functional SERCA2a-DWORF interactions. This work advances our mechanistic understanding of the regulation of SERCA2a by small transmembrane proteins and sets the stage for future therapeutic development in heart failure research.

## Introduction

Sarcoplasmic reticulum (SR) calcium ATPase (SERCA) regulates cytosolic Ca^2+^ levels within muscle cells (1). SERCA is responsible for pumping the majority of Ca^2+^ from the cardiomyocyte sarcoplasm into the SR lumen to enable muscle relaxation (diastole) after each muscle contraction (systole),(2) using energy derived from ATP hydrolysis (3). There are three major domains on the cytoplasmic face of SERCA. The phosphorylation (P) and nucleotide-binding (N) domains form the catalytic site, while the actuator (A) domain is involved in the transduction of conformational changes required for the Ca-uptake function involving the transmembrane (TM) domain (4). Of the twelve known SERCA isoforms expressed in muscle and non-muscle cells (5), SERCA2a is the principal isoform expressed in cardiac myocytes. Reduced SERCA2a expression and/or activity has been identified as a causative factor contributing to the development of heart failure (HF) in humans (6). Specifically, elevated cytoplasmic Ca^2+^ levels in cardiomyocytes cause impaired muscle relaxation during diastole and extended duration of systole, and these effects are strongly correlated with the pathophysiology of HF (7,8). Given that cardiovascular disease is now a major health concern worldwide (9), SERCA2a represents an attractive target to directly mitigate cardiomyocyte deficiencies associated with pathological states.

SERCA2a is regulated by several homologous single-pass, helical TM proteins, including phospholamban (PLB) (10,11), a 52-residue protein that is co-expressed with SERCA2a in ventricular myocytes. Unphosphorylated PLB inhibits SERCA2a, but inhibition is relieved by either micromolar Ca^2+^ (systolic) or β-adrenergic stimulation of PLB phosphorylation (12). Hereditary mutations in PLB increase inhibitory potency and are associated with human HF (10). Therefore, gene and small-molecule therapies to increase SERCA2a activity, either by activating SERCA2a directly or by relieving inhibition by PLB (10,13,14) are of strong interest to the research and medical community. One gene therapy approach has been to express non-inhibitory PLB mutants, evaluated by their ability to displace the inhibitory wild type PLB, as measured by FRET between a donor fluorophore on SERCA2a and an acceptor fluorophore on PLB (15–17).

All long-known endogenous membrane protein regulators of SERCA (*e.g*., PLB, sarcolipin) are inhibitory, but a SERCA-activating peptide termed dwarf open reading frame (DWORF) has recently been identified (18). DWORF activates SERCA2a in cardiac myocytes, where PLB is also present, leading to the hypothesis that DWORF activates SERCA2a indirectly by displacing PLB but has no direct effect on SERCA2a function (18). In the present study, we have investigated this mechanism by using fluorescence lifetime (FLT) detection of FRET, to determine quantitatively the interactions of PLB and DWORF (separately and in combination) with SERCA2a, and correlate with functional assays of SERCA activity on the same samples.(16,19). With this approach, we find that DWORF increases the Ca-ATPase activity of SERCA not only by displacing the inhibitory PLB, but also through direct interactions with SERCA2a in the absence of PLB. Using site-directed mutagenesis,, we identified two DWORF residues, P15 and W22 (in the human isoform), as essential for activation of SERCA2a. These findings may provide key insights needed to develop effective therapies for heart failure.

## Results

### Expression, localization, and FRET measurements of SERCA2a-DWORF biosensor

To study the SERCA2a-DWORF interaction in live cells, we developed a FRET-based biosensor by fusing GFP and tagRFP to the N-termini of SERCA2a and DWORF, respectively. In this biosensor, GFP-SERCA (the FRET donor) is stably expressed, while RFP-DWORF (the FRET acceptor) is transiently expressed to allow controlling the expression levels of the acceptor. Fluorescence microscopy confirmed that both GFP-SERCA2a and RFP-DWORF were expressed and co-localized in the ER (Fig. 1A). Fluorescence measurements of the donor lifetime in the presence of the acceptor (GFP-SERCA2a + RFP-DWORF) showed a significantly faster decay, compared to the donor only (GFP-SERCA2a) samples (Fig. 1B). FRET efficiency (E) and the fraction of donor molecules in proximity to acceptor molecules (X_DA_) were calculated using multiexponential FLT data analysis (see Methods). Due to the R^−6^ dependence of FRET on the donor-acceptor distance R, X_DA_ is a reliable measurement of the fraction of donor-labeled SERCA population that has bound acceptor-labeled PLB or DWORF (15). We found that increasing RFP-DWORF expression results in increasing FRET (E) and X_DA_ (Fig. 1C-D), indicating that RFP-DWORF and GFP-SERCA2a are expressed and bound to each other in live cells.

**Figure 1.**
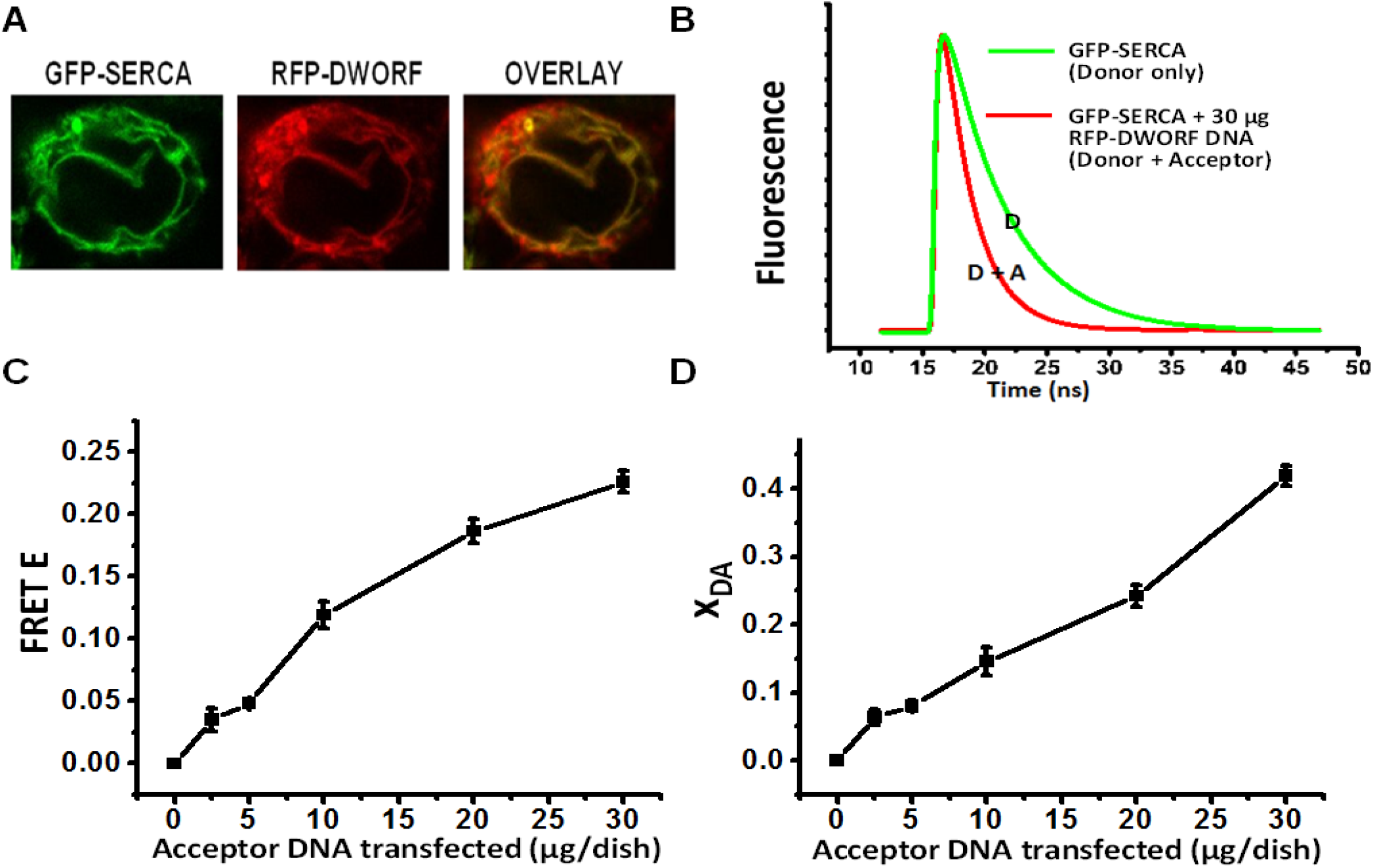
RFP-DWORF binding to GFP-SERCA2a in live cells. (A) Confocal microscopy images of GFP-SERCA (donor) and RFP-DWORF (acceptor) fluorescence signals. (B) Fluorescence lifetime (FLT) traces measured from HEK293 cells stably expressing the donor without (green, F_D_(t) or with (red, F_D+A_(t)) transient expression of the acceptor. FLT data was analyzed as described in Methods to determine lifetimes (τ_D_ and τ_D+A_) and mole fractions X_D_ and X_DA_. (C) FRET efficiency E (1-τ_D+A_/τ_D_) vs. μg of acceptor DNA seeded per dish. (D) The fraction of SERCA2a containing bound DWORF (X_DA_).

### DWORF competes with PLB for binding to SERCA2a

We transfected increasing amounts of a plasmid carrying unlabeled DWORF into a stable cell line expressing GFP-SERCA2a (donor) and RFP-PLB (acceptor) (Fig. 2A). In the absence of DWORF, FRET (E) between labeled SERCA and PLB was 0.13 ± 0.04, and X_DA_ = 0.30 ± 0.03 (Fig. 2B-C). Addition of unlabeled DWORF results in decreased FRET efficiency, indicating a decrease in the interaction between SERCA2a and PLB, most likely due to a decrease in the fraction of SERCA containing bound PLB (X_DA_) (Fig. 2). We conclude that DWORF competes effectively with PLB binding to SERCA2a.

**Figure 2.**
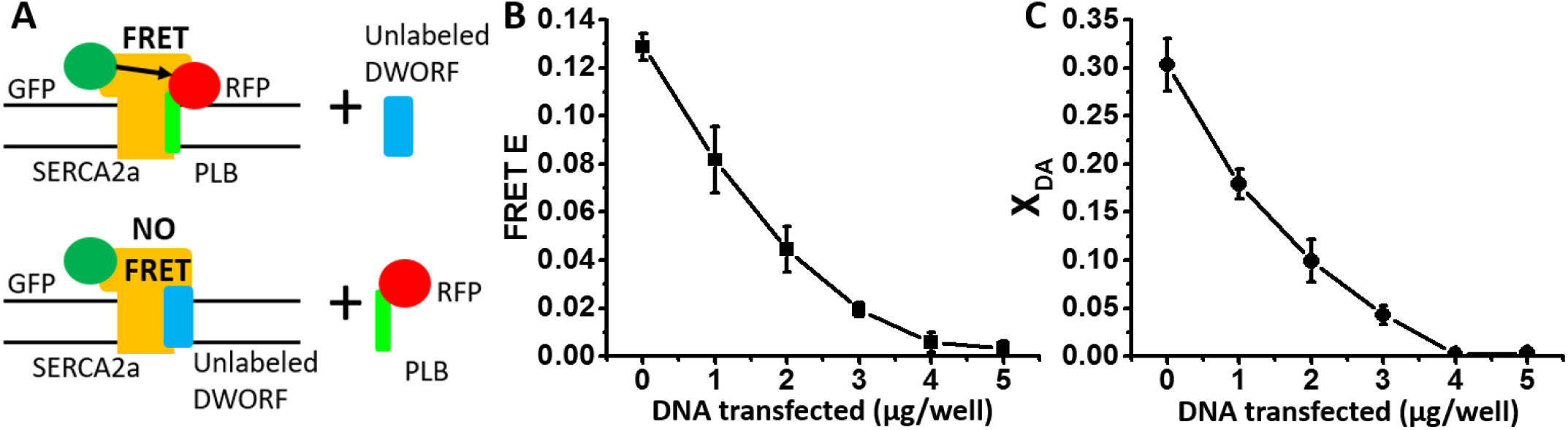
DWORF competes with PLB for SERCA binding. (A) Schematic illustration of the tested hypothesis, of competition between unlabeled DWORF and RFP-PLB for binding to GFP-SERCA2a. (B) FRET efficiency (E) vs. μg/well of DWORF DNA transfected into 6-well plates stably expressing GFP-SERCA2a and RFP-PLB. (C) Mole fraction (X_DA_) of GFP-SERCA (donor) bound to RFP-PLB (acceptor), for the same samples as in B. n=3

### TM domain of DWORF is sufficient for functional interaction with SERCA2a

DWORF consists of a 23-residue TM (9) domain and an 11-residue cytoplasmic domain that forms a short helix on the cytoplasmic side of the SR membrane (18). To map which of these domains is responsible for functional interaction with SERCA2a, we engineered a truncated DWORF mutant containing only the TM domain. The RFP-labeled TM domain of DWORF (RFP-TM-DWORF) was transiently transfected into HEK293 cells stably expressing GFP-SERCA2a. With increasing RFP-TM-DWORF expression, we observed increases in both FRET (E) and X_DA_ values, similar to those observed with full-length DWORF (Fig. 3A-B). This suggests that the TM domain of DWORF interacts with SERCA similarly to full-length DWORF.

**Figure 3.**
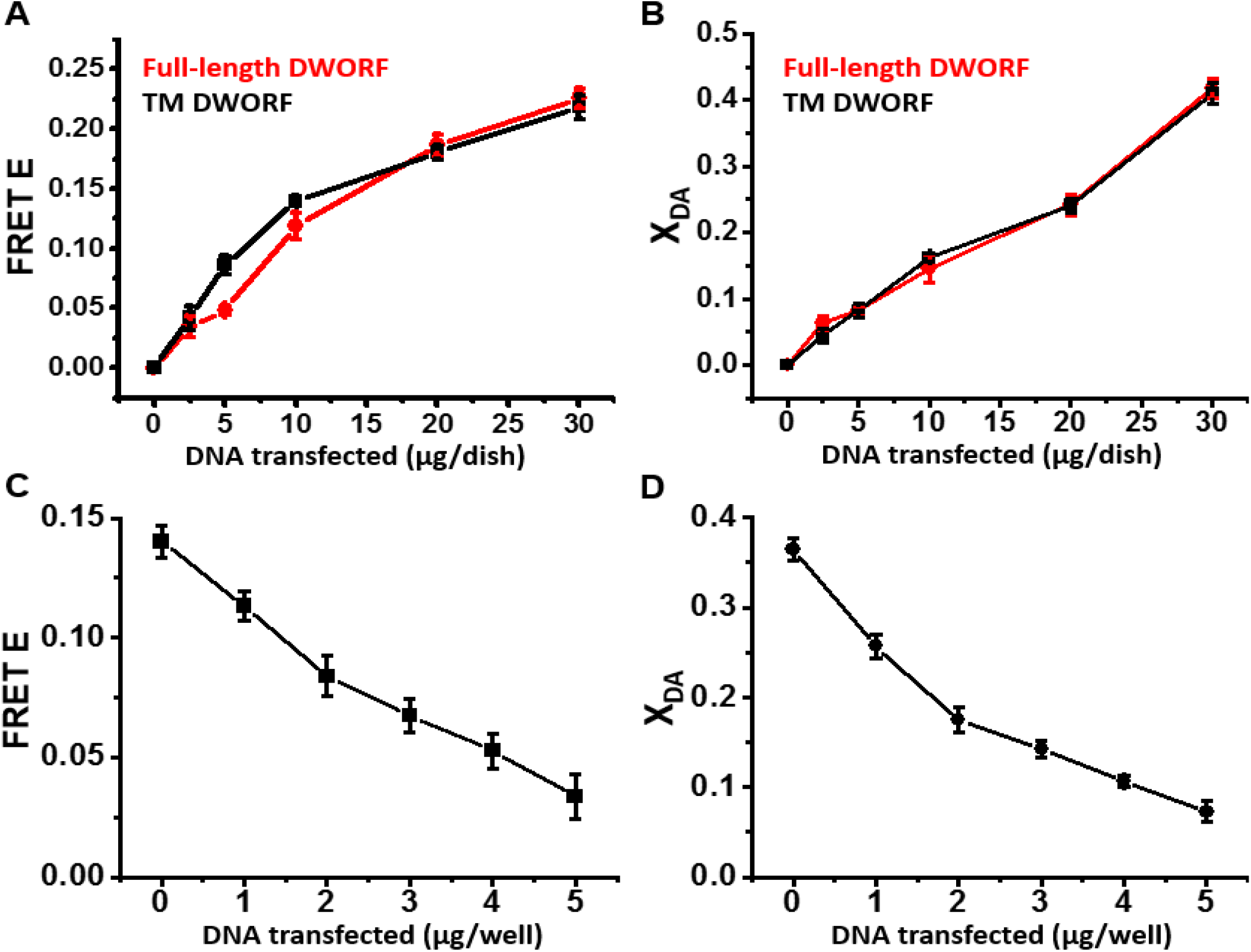
FRET-detected interaction of SERCA with the isolated TM domain of DWORF is similar to that with full-length DWORF. (A) FRET efficiency (E) in HEK293 cells stably expressing GFP-SERCA2a, vs. μg/dish of DNA transiently transfected into HEK293 cells expressing RFP-TM-DWORF (black) or full-length RFP-DWORF (red). (B) Fraction X_DA_ of donors (GFP) transferring energy (bound) to acceptor (RFP) determined for samples in panel A. (C) FRET efficiency E for increasing amounts of TM-DWORF DNA transfected into a stable cell line expressing GFP-SERCA2a and RFP-PLB. (D) FRET efficiency plotted vs. DNA transfected per well. D. Fraction X_DA_ of donors transferring energy (bound) to acceptors determined for samples in C.

Next, we performed competition experiments using an unlabeled TM-DWORF construct, to quantify binding of the TM-DWORF to GFP-SERCA2a in the presence of RFP-PLB. As in experiments using full length DWORF (Fig. 2C), we observed a decrease in the FRET (E) between labeled SERCA2a and PLB (Fig. 3C). X_DA_, the fraction of GFP-SERCA2a transferring energy to RFP-PLB was decreased (Fig. 3D). We conclude that the TM domain of DWORF is sufficient to compete with PLB for binding to SERCA2a.

### P15 and W22 residues in DWORF (human isoform) are essential for full extent of binding to SERCA2a

To determine which residues within the TM domain of DWORF are important for complex formation with SERCA2a, DWORF mutants were selected for testing based on evolutionary conservation (Fig. 4A). Initially, we tested the effects on single mutations on the ability of fluorescently labeled DWORF constructs to bind to labeled SERCA2a and participate in FRET. However, we did not observe significant differences in the binding affinities of single-mutation DWORF constructs for SERCA2a, as compared to the wild-type controls (Fig. 4C). Given that the protein-protein interface between TM domains of the SERCA-DWORF complex may involve interactions involving multiple DWORF residues, we set out to test whether a combination of two mutations is sufficient to ablate complex formation. We found that one combination, a double mutant in which P15 and W22 are both substituted with alanine, showed significant decreases in both FRET efficiency and X_DA_ (Fig. 4D-E), suggesting that this mutant DWORF binds SERCA2a with a significantly lower affinity than the WT-DWORF controls. We also tested whether the unlabeled P15A/W22A DWORF mutant can effectively compete for binding to GFP-SERCA in the presence of RFP-PLB and found that there is little to no competitive binding. (Fig. 4F). Based on the DWORF TM sequence, which is predicted to form a single helix, both P15 and W22 residues are on the same face of the helix (Fig. 4B), suggesting that this is the protein-protein interface between the TM domains of SERCA2a and DWORF.

**Figure 4.**
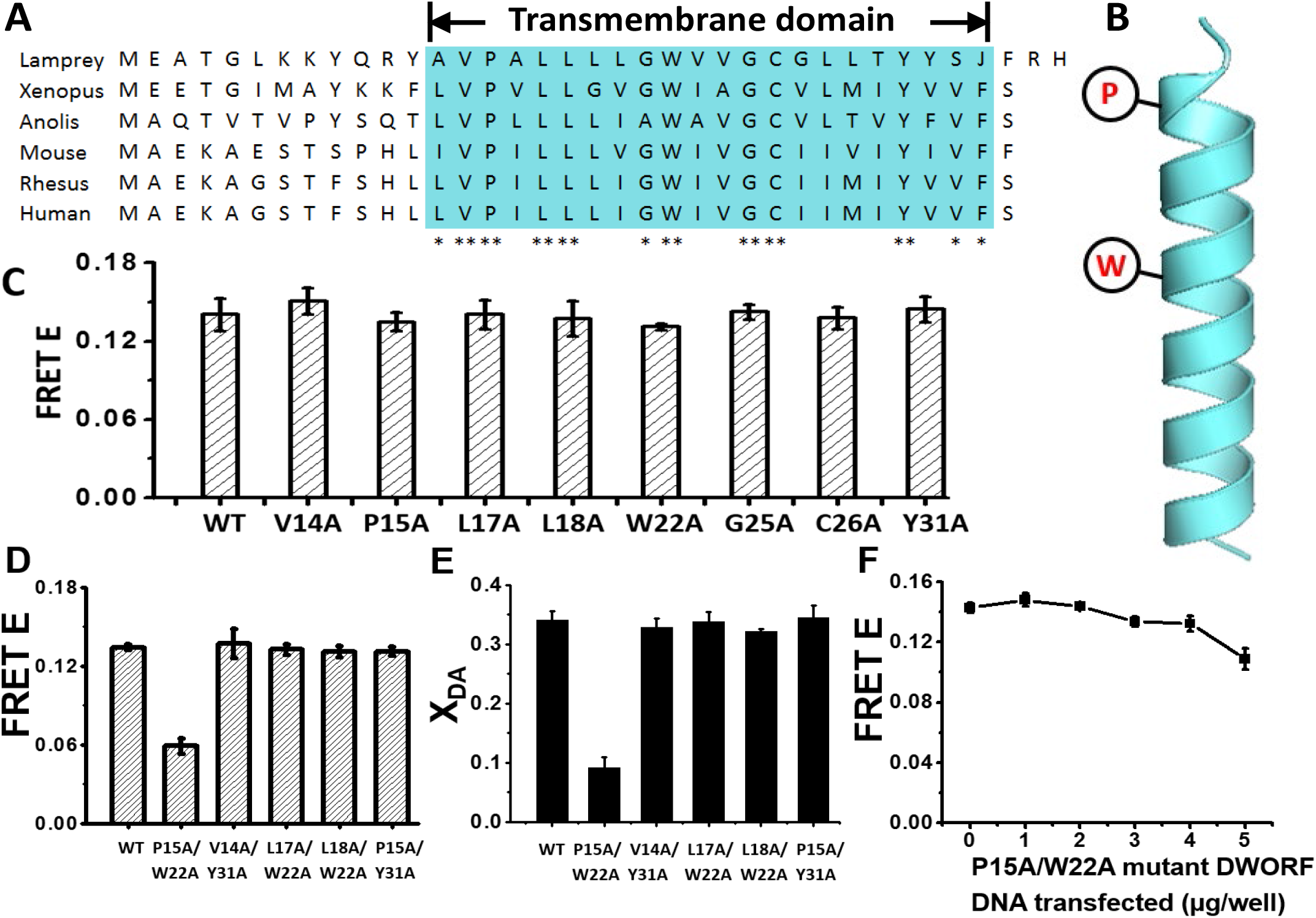
P15 and W22 residues of DWORF (human isoform) are essential for the full extent of DWORF-SERCA2a complex formation. (A) DWORF amino acid sequence alignment [18]. Blue region denotes the putative TM domain. ** indicates a residue conserved among all species shown; * indicates a residue partially conserved. (B) Structure of DWORF TM domain, with P15 and W22 indicated in red. (C) FRET efficiency of RFP-DWORF single-mutant constructs expressed with GFP-SERCA2a. 5μg/well of each mutant DWORF DNA was transfected into a stable cell line expressing GFP-SERCA2a. (D) FRET efficiency of RFP-DWORF double-mutant constructs expressed under similar conditions. 5μg/well of each double mutant DWORF DNA was transfected into a stable cell line expressing GFP-SERCA2a. (E) Fraction X_DA_ of donors transferring energy (bound) to acceptor determined for samples in D. (F) FRET efficiency E vs μg/well of RFP-DWORF (P15A/W22A) transfected. n=3.

### DWORFactivates SERCA2a in the absence of PLB

Based on FRET measurements, it is clear that DWORF competes effectively with PLB for binding to SERCA2a (Fig. 2), and this competition has been shown to reverse PLB inhibition of SERCA2a (18). FRET shows that DWORF also binds to SERCA in the absence of PLB (Fig. 1). To test the hypothesis that DWORF activates SERCA2a directly, in the absence of PLB, we performed Ca-ATPase assays in HEK293 cells overexpressing GFP-SERCA2a and varying amounts of DWORF, in the absence of PLB (Fig. 5). Surprisingly, we found that *DWORF activates SERCA2a directly* at sub-saturating [Ca^2+^], increasing the apparent Ca affinity of SERCA2a (Fig. 5A). Similar results were obtained with RFP-DWORF (Fig. S1). As a control, we tested the effects of the double mutant P15A/W22A DWORF, which displays low affinity toward SERCA2a in our FRET assays, and found that this construct has no effect on Ca-ATPase function (Fig. 5B).

**Figure 5.**
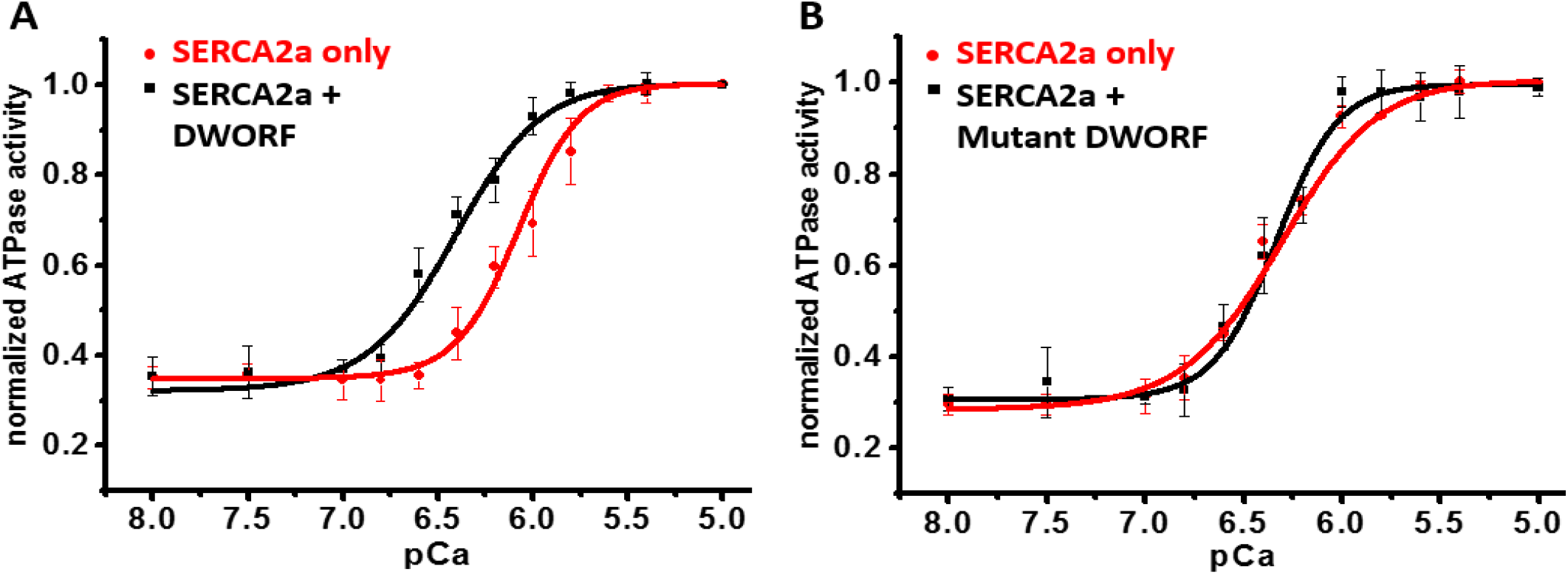
DWORF activates SERCA2a in the absence of PLB. (A) Ca-ATPase activity measured in homogenates of cells expressing GFP-SERCA2a and unlabeled DWORF (black, pK_Ca_ = 6.36 ± 0.04) or GFP-SERCA2a only (red, pK_Ca_ = 6.08 ± 0.03). (B) Ca-ATPase activity measured in homogenates of cells expressing GFP-SERCA2a and unlabeled DWORF-P15/W22 (black, pK_Ca_ = 6.36 ± 0.04) or GFP-SERCA2a only (red, pK_Ca_ = 6.08 ± 0.03). GFP-SERCA2a was stably expressed in HEK293 cells, then 20 μg of DNA per dish (for unlabeled DWORF or unlabeled mutant DWORF) was transiently transfected into the cells. Ca-ATPase assays were performed using cell homogenates as described in Methods. Activity was normalized to the control (SERCA2a only), since DWORF had no significant effect on V_max_. (n=3)

## Discussion

By combining structural (FRET) and functional (Ca-ATPase) measurements, the present study reveals a new role for DWORF in calcium regulation. In our model, DWORF binds to SERCA2a, displacing PLB, and activates SERCA via two distinct mechanisms: (1) DWORF displaces the inhibitory PLB and (2) DWORF activates SERCA directly; above the level of SERCA in the absence of both DWORF and PLB. Residues P15 and W22 in the TM domain of DWORF are required for DWORF-SERCA binding.

The causes of HF are complex, but growing evidence indicates that central factors include increased basal cytosolic [Ca^2+^], correlated with decreased [Ca^2+^] in the SR lumen. Since SERCA2a is the major determinant of cytosolic [Ca^2+^], increasing SERCA2a activity is a potential therapy for a myriad of chronic cardiac pathologies (20). In support of this hypothesis, DWORF enhances SERCA2a activity and contractility in a HF mouse model (21). Here we study this mechanism through direct detection of protein-protein complex formation, using FRET biosensors based on DWORF, SERCA2a, and PLB. We observed that DWORF binds directly to SERCA2a (Fig. 1) and competes effectively with PLB for SERCA2a binding (Fig. 2). This is consistent with previous *in vivo* observations (18) and fluorescence-based measurements showing that SERCA has a higher apparent affinity for DWORF as compared to PLB (21). DWORF is reported to interact with SERCA2a to form a heterodimer (22). We observed that DWORF activates SERCA2a directly, in the absence of PLB, by enhancing the Ca-ATPase apparent calcium affinity (Fig 5A). This direct activation of SERCA2a by DWORF was not detected in previous measurements (22). This difference may be due to differences in SERCA2a-DWORF stoichiometry, cell lines used, and/or the presence of protein tags. Given that we measured Ca-ATPase activity using an unlabeled construct at the same SERCA2a-DWORF stoichiometry as in our structure-based assays, we propose that these findings reveal a new role for DWORF, in which DWORF functions as a *bona fide* activator of SERCA2a.

We have shown that the TM domain of DWORF is sufficient for complex formation with SERCA2a and subsequent activation (Fig. 3) (Fig. S2). We identified two resides, P15 and W22, that are important for this interaction (Fig. 4). When P15 and W22 are both mutated to A, the fraction of SERCA2a containing bound DWORF decreases by a factor of four (Fig. 5B), indicating that this double mutant fails to activate SERCA2a primarily due to decreased binding. However, the double mutant did not completely eliminate FRET or X_DA_, the fraction of SERCA containing bound DWORF (Fig. 4D-E), indicating that the double mutation eliminates activation by bound DWORF. Future structural studies (*e.g*., crystallography and cryo-EM) will reveal the complete protein-protein interface, and further develop the mechanistic model of DWORF’s function.

## Conclusion

The current study demonstrates that DWORF binds to SERCA2a, competing with PLB (Fig. 2). The binding of DWORF to SERCA2a depends entirely on the transmembrane (TM) domain of DWORF (Fig. 3), and mutation of TM residues P15 and W22 to A greatly decreases the binding of DWORF to SERCA2a (Fig. 4). These results are consistent with the previous proposal that DWORF activates SERCA2a by displacing inhibitory PLB from SERCA (18). However, we find that ***DWORF also activates SERCA2a directly*** in the absence of PLB, while the double mutant has no effect (Fig. 5). These results will help to inform future efforts in therapeutic design; a better understanding of the regulation of SERCA2a by small transmembrane proteins is essential for development of new classes of therapeutics in heart failure research.

## Experimental procedures

### Molecular biology

The EGFP gene was fused to that of the N-terminus of human SERCA2a gene plus a 5-residue linker, as described previously (16,23). The tagRFP gene was similarly fused to the N-terminus of the human DWORF or PLB gene. The GFP-SERCA2a construct was cut and pasted with NheI and NotI cutting sites into a pcDNA3.1 vector with ampicillin resistance. The genes for RFP-DWORF, RFP-TM domain of DWORF (RFP-TMDWORF), RFP-PLB, RFP-DWORF, and unlabeled DWORF were separately cut and pasted with NheI and NotI into a pEGFPc1 vector with Kanamycin resistance. RFP-TMDWORF fragment with NheI and NotI cutting site was ordered through the University of Minnesota Genomics Center (UMGC), then inserted into the pEGFPc1 vector. Mutagenesis of DWORF was preformed using Q5 Site-Directed Mutagenesis Kit (New England BioLabs, MA).

### Cell Culture and transfections

HEK293 cells were transfected using Lipofectamine 3000 (Thermo Fisher). SERCA2a constructs were expressed using the mammalian expression vector pcDNA3.1 with G418 resistance. DWORF and PLB constructs were expressed using mammalian expression vector peGFPc1 with puromycin resistance. 24 hours before transfection, HEK293 cells were plated at 0.4 million cells per well in 6-well plates, or 2 million cells per dish in 10-centimeter dishes. Different amounts of DNA were transfected using Lipofectamine 3000 (Thermo Fisher). For FRET lifetime detection, cells were be harvested at 1 million cells per mL in PBS 48 hours after transfection. To select the stable cell line, two days after transfection, 2.0 mg/mL puromycin or 500 μg/mL G418 antibiotic was added to the growth medium. Seven days after antibiotic selection, the remaining cells were enriched by fluorescence-activated cell sorting (FACS). After three weeks in culture, there were approximately 100 million cells, generating a stable clone expressing the biosensor at high levels. The stable cell line was maintained using F17 medium (Sigma) + (200nM/mL) GlutaMAX + 2.0 mg/mL puromycin or 500 μg/mL G418.

### Preparation of Cell Homogenates

Homogenate preparation was performed as previously reported (24). Briefly, cells were centrifuged at 300xg, washed three times in phosphate buffer solution (PBS, with no magnesium or calcium added, Thermo Scientific), and resuspended in homogenization buffer (0.5mM MgCl_2_, 10mM Tris-HCL pH 7.5, DNase I and protease inhibitor) at ten million cells/mL. Cells were then incubated on ice for ten minutes, and broken with the Tissumizer (Tekmar SDT-1810) with three 30-second bursts. After each burst, a 5 minute incubation on ice was performed. Homogenization was confirmed with a microscope. After the homogenization, 2x sucrose buffer (1mM MOPS, 500mM sucrose, and protease inhibitor) was added for a final cell concentration of 2mg total protein per mL.

### NADH-enzyme coupled ATPase activity assay

Functional assays were performed using homogenate preparations expressing either the SERCA2a-DWORF biosensor or the appropriate donor-only control. The protocol of ATPase activity assays similar as previously described (24). An enzyme-coupled, NADH-linked ATPase assay was used to measure SERCA2a Ca-ATPase activity in 96-well microplates. Each well contained assay mix (50 mM MOPS pH 7.0, 100 mM KCl, 5 mM MgCl_2_, 1 mM EGTA, 0.2 mM NADH, 1 mM phosphoenol pyruvate, 10 IU/mL of pyruvate kinase, 10 IU/mL of lactate dehydrogenase, 1 μM of the calcium ionophore A23187 from Sigma [St. Louis, MO]), and added CaCl_2_ to set the free [Ca^2+^] to the desired values. Before starting the assay, 4 mg/mL of cell homogenate, CaCl_2_, compound, and assay mix were incubated for 20 min. The assay was then started by adding ATP to a final concentration of 5 mM (200 mL total assay volume), and absorbance was measured at 340 nm in a SpectraMax Plus microplate spectrophotometer (Molecular Devices, Sunnyvale, CA). Data points were fitted with V = V_0_ + V_max_/(1 + 10^−n[pKCa – pCa]^), to determine pK_Ca_, the p_Ca_ value for half-maximal activation by Ca. PLB causes an decrease in pK_Ca_ (increase in [Ca^2+^] required for SERCA activation), and activation of SERCA corresponds to an increase in pK_Ca_.

### FLT measurement and data analysis

FLT measurements were performed by time-correlated single photon counting (TCSPC), as previously described (25). HEK293 cells expressing fluorescent biosensors were harvested at 1 million cells per mL in PBS. TCSPC experiments were performed on samples containing water (instrument response function, IRF), untransfected HEK293 (background), and HEK293 expressing donor only (GFP-SERCA2a) and donor plus acceptor (RFP-DWORF or RFP-PLB). TCSPC experiments were performed following excitation at 481 nm using a subnanosecond pulsed diode laser, selecting emitted light using a 540 ± 20 nm filter (Semrock, NY), and detecting with a PMH-100 photomultiplier (Becker-Hickl).

FRET-labeled samples were analyzed as described in our previous publications (12,25–27). Data was first analyzed assuming a single FLT τ_D_, for the sample containing donor only (GFP-SERCA2a), and τ_D+A_ for the sample containing both donor and acceptor. FRET efficiency E was calculated as E = 1 – (τ_D+A_/τ_D_). The data for the D+A sample was further analyzed according to F_D+A_(t) = X_D_F_D_(t) + X_DA_F_DA_(t), where F_DA_(t) is the fluorescence of the donor-acceptor complex, and X_DA_ is the mole fraction of donor (GFP-SERCA2a) containing bound acceptor (RFP-PLB).

### Statistical analyses

Analysis of two group comparisons was done by Student’s t-test (*P < 0.05). Data are presented as mean ± standard error of the mean (SEM), and all statistical values were calculated from a minimum of three separate experiments.

## Data availability

All data discussed are presented within the article.

## Acknowledgements

Ji Li, Prachi Bawaskar, Mark Rustad, Tory M. Schaaf, and J. Michael Autry provided helpful technical advice and discussion, Destiny Ziebol assisted with manuscript preparation, and Sarah Anderson provided administrative support. Spectroscopy was performed at the UMN Biophysical Technology Center.

## Funding and additional information

This work was supported in part by NIH grants R01HL139065 (to DDT and RLC; formerly R01GM027906), and R37AG026160 (to DDT).

## Author Contributions

AL, RLC, and DDT designed the research. AL, DRS, SLY and EK prepared samples, performed experiments, and analyzed the data. AL, DRS, RLC and DDT wrote the paper.

## Supporting information

**Figure S1.**
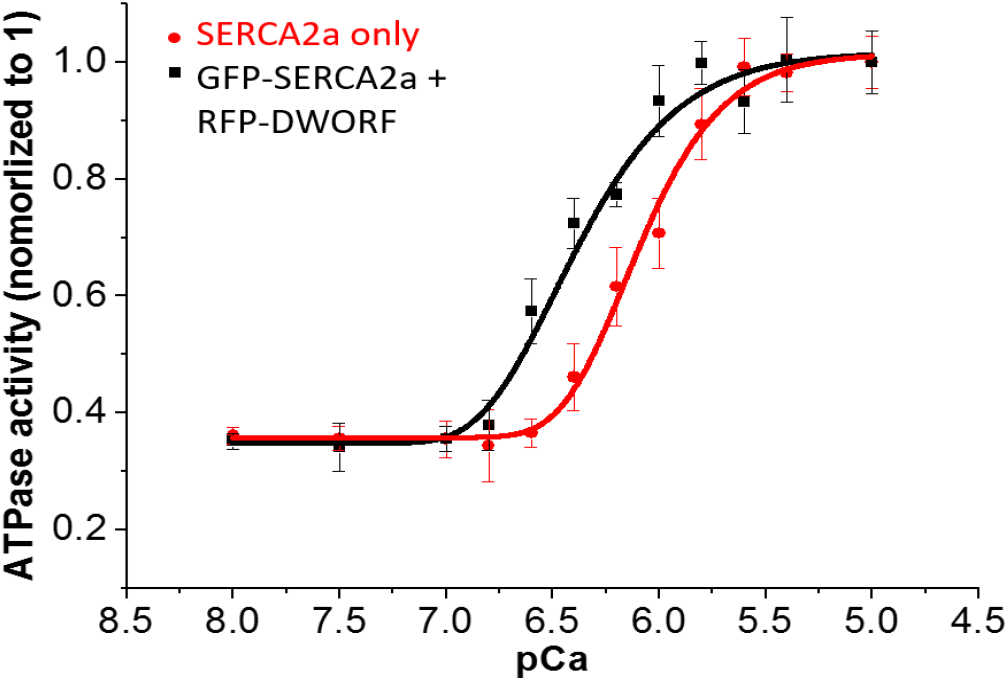
RFP-DWORF activates SERCA2a directly in the absence of PLB. Transient transfections of 5μg of RFP-DWORF into HEK293 cells stably expressing GFP-SERCA2a. 48 hours after transfection, cells was harvested at 10 million per mL in homogenization buffer (0.5mM MgCl2, 10mM Tris-HCL ph 7.5, DNase I and protease inhibitor). Cells was then lysed by Tissumizer (Tekmar SDT-1810) and prepared in 96well plate for ATPase assay. n=3

**Figure S2.**
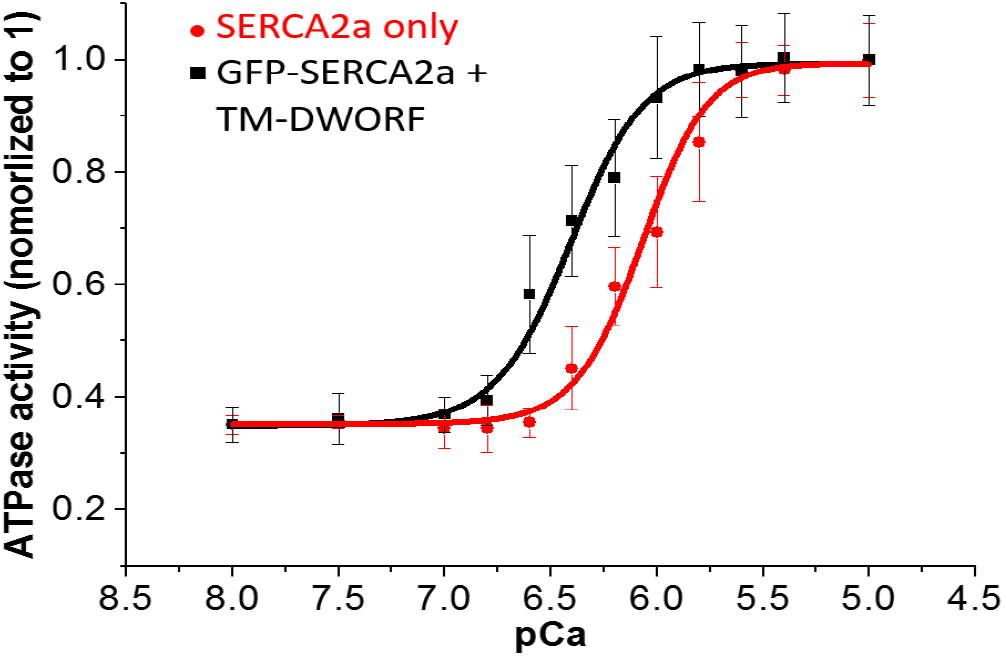
TM-DWORF can activate SERCA2a directly in the absence of PLB. Transient transfections of 5μg of TM-DWORF into HEK293 cells stably expressing GFP-SERCA2a. 48 hours after transfection, cells was harvested at 10 million per mL in homogenization buffer (0.5mM MgCl2, 10mM Tris-HCL ph 7.5, DNase I and protease inhibitor). Cells was then lysed by Tissumizer (Tekmar SDT-1810) and prepared in 96well plate for ATPase assay. n=3

**Figure S3.**
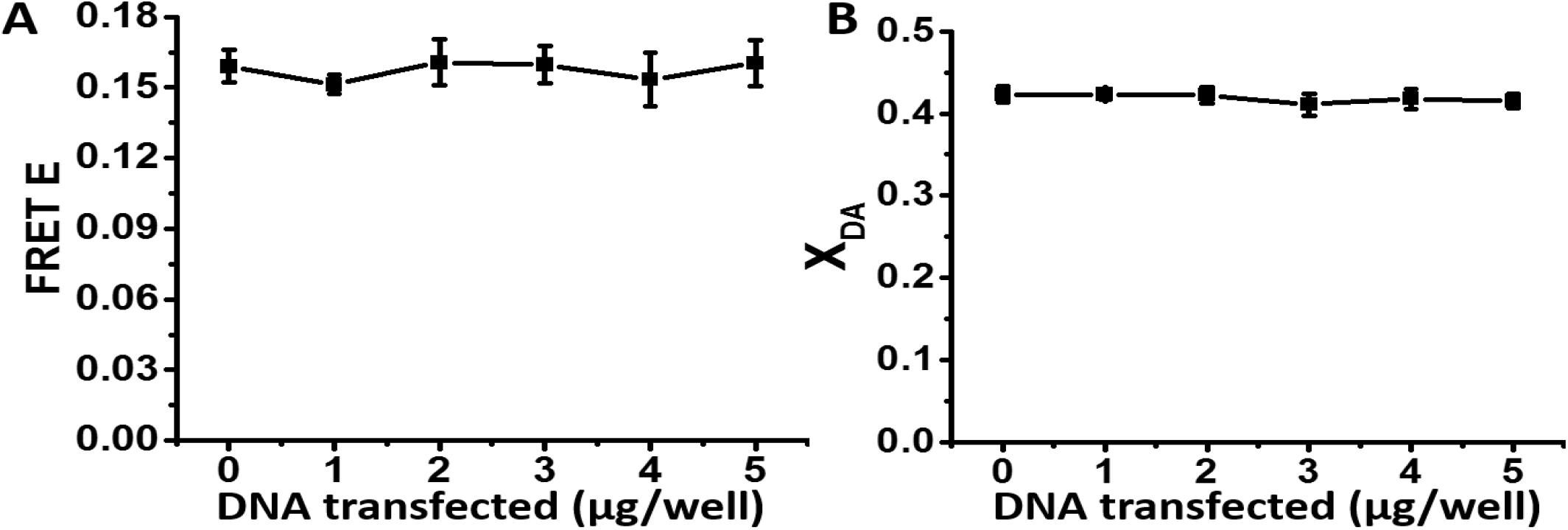
Unlabeled PLB does not compete with labeled DWORF. (A) FRET E to measure competition between PLB and DWORF. Different amounts of unlabeled PLB DNA and 3μg of RFP-DWORF DNA were transiently transfected into HEK293 cells stably expressing GFP-SERCA2a. 48 hours after transfection, cells were harvested at 1 million per mL in PBS for FLT measurements of FRET. (B) Fraction of donors transferring energy to acceptor (determined by fitting time-resolved fluroescence data to. n=3

## References

1. Asahi, M., Otsu, K., Nakayama, H., Hikoso, S., Takeda, T., Gramolini, A. O., Trivieri, M. G., Oudit, G. Y., Morita, T., Kusakari, Y., Hirano, S., Hongo, K., Hirotani, S., Yamaguchi, O., Peterson, A., Backx, P. H., Kurihara, S., Hori, M., and MacLennan, D. H. (2004) Cardiac-specific overexpression of sarcolipin inhibits sarco(endo)plasmic reticulum Ca2+ ATPase (SERCA2a) activity and impairs cardiac function in mice. Proc Natl Acad Sci U S A 101, 9199–9204

2. Bers, D. M. (2002) Cardiac excitation-contraction coupling. Nature 415, 198–205

3. Thomas, D. D., and Karon, B. S. (1994) Temperature dependence of molecular dynamics and calcium ATPase activity in sarcoplasmic reticulum. in The Temperature Adaptation of Biological Membranes. (Cossins, A. R. ed.), Portland Press, London,. pp 1–12

4. Smeazzetto, S., Saponaro, A., Young, H. S., Moncelli, M. R., and Thiel, G. (2013) Structure-function relation of phospholamban: modulation of channel activity as a potential regulator of SERCA activity. PLoS One 8, e52744

5. Periasamy, M., and Kalyanasundaram, A. (2007) SERCA pump isoforms: their role in calcium transport and disease. Muscle Nerve 35, 430–442

6. Dode, L., Andersen, J. P., Leslie, N., Dhitavat, J., Vilsen, B., and Hovnanian, A. (2003) Dissection of the functional differences between sarco(endo)plasmic reticulum Ca2+-ATPase (SERCA) 1 and 2 isoforms and characterization of Darier disease (SERCA2) mutants by steady-state and transient kinetic analyses. J Biol Chem 278, 47877–47889

7. Smith, K. A., Ayon, R. J., Tang, H., Makino, A., and Yuan, J. X. (2016) Calcium-Sensing Receptor Regulates Cytosolic [Ca (2+)] and Plays a Major Role in the Development of Pulmonary Hypertension. Front Physiol 7, 517

8. Kranias, E. G., and Hajjar, R. J. (2012) Modulation of cardiac contractility by the phospholamban/SERCA2a regulatome. Circ Res 110, 1646–1660

9. Mozaffarian, D., Benjamin, E. J., Go, A. S., Arnett, D. K., Blaha, M. J., Cushman, M., de Ferranti, S., Despres, J. P., Fullerton, H. J., Howard, V. J., Huffman, M. D., Judd, S. E., Kissela, B. M., Lackland, D. T., Lichtman, J. H., Lisabeth, L. D., Liu, S., Mackey, R. H., Matchar, D. B., McGuire, D. K., Mohler, E. R., 3rd, Moy, C. S., Muntner, P., Mussolino, M. E., Nasir, K., Neumar, R. W., Nichol, G., Palaniappan, L., Pandey, D. K., Reeves, M. J., Rodriguez, C. J., Sorlie, P. D., Stein, J., Towfighi, A., Turan, T. N., Virani, S. S., Willey, J. Z., Woo, D., Yeh, R. W., Turner, M. B., American Heart Association Statistics, C., and Stroke Statistics, S. (2015) Heart disease and stroke statistics--2015 update: a report from the American Heart Association. Circulation 131, e29–322

10. Ceholski, D. K., Turnbull, I. C., Kong, C. W., Koplev, S., Mayourian, J., Gorski, P. A., Stillitano, F., Skodras, A. A., Nonnenmacher, M., Cohen, N., Bjorkegren, J. L. M., Stroik, D. R., Cornea, R. L., Thomas, D. D., Li, R. A., Costa, K. D., and Hajjar, R. J. (2018) Functional and transcriptomic insights into pathogenesis of R9C phospholamban mutation using human induced pluripotent stem cell-derived cardiomyocytes. J Mol Cell Cardiol 119, 147–154

11. MacLennan, D. H., and Kranias, E. G. (2003) Phospholamban: a crucial regulator of cardiac contractility. Nat Rev Mol Cell Biol 4, 566–577

12. Dong, X., and Thomas, D. D. (2014) Time-resolved FRET reveals the structural mechanism of SERCA-PLB regulation. Biochemical and biophysical research communications 449, 196–201

13. Stammers, A. N., Susser, S. E., Hamm, N. C., Hlynsky, M. W., Kimber, D. E., Kehler, D. S., and Duhamel, T. A. (2015) The regulation of sarco(endo)plasmic reticulum calcium-ATPases (SERCA). Can J Physiol Pharmacol 93, 843–854

14. Schmitt, J. P., Kamisago, M., Asahi, M., Li, G. H., Ahmad, F., Mende, U., Kranias, E. G., MacLennan, D. H., Seidman, J. G., and Seidman, C. E. (2003) Dilated cardiomyopathy and heart failure caused by a mutation in phospholamban. Science 299, 1410–1413

15. Stroik, D. R., Yuen, S. L., Janicek, K. A., Schaaf, T. M., Li, J., Ceholski, D. K., Hajjar, R. J., Cornea, R. L., and Thomas, D. D. (2018) Targeting protein-protein interactions for therapeutic discovery via FRET-based high-throughput screening in living cells. Sci Rep 8, 12560

16. Stroik, D. R., Ceholski, D. K., Bidwell, P. A., Mleczko, J., Thanel, P. F., Kamdar, F., Autry, J. M., Cornea, R. L., and Thomas, D. D. (2020) Viral expression of a SERCA2a-activating PLB mutant improves calcium cycling and synchronicity in dilated cardiomyopathic hiPSC-CMs. J Mol Cell Cardiol 138, 59–65

17. Gruber, S. J., Haydon, S., and Thomas, D. D. (2012) Phospholamban mutants compete with wild type for SERCA binding in living cells. Biochem Biophys Res Commun 420, 236–240

18. Nelson, B. R., Makarewich, C. A., Anderson, D. M., Winders, B. R., Troupes, C. D., Wu, F., Reese, A. L., McAnally, J. R., Chen, X., Kavalali, E. T., Cannon, S. C., Houser, S. R., Bassel-Duby, R., and Olson, E. N. (2016) A peptide encoded by a transcript annotated as long noncoding RNA enhances SERCA activity in muscle. Science 351, 271–275

19. Cornea, R. L., Gruber, S. J., Lockamy, E. L., Muretta, J. M., Jin, D., Chen, J., Dahl, R., Bartfai, T., Zsebo, K. M., Gillispie, G. D., and Thomas, D. D. (2013) High-throughput FRET assay yields allosteric SERCA activators. J Biomol Screen 18, 97–107

20. Zhihao, L., Jingyu, N., Lan, L., Michael, S., Rui, G., Xiyun, B., Xiaozhi, L., and Guanwei, F. (2020) SERCA2a: a key protein in the Ca(2+) cycle of the heart failure. Heart Fail Rev 25, 523–535

21. Makarewich, C. A., Munir, A. Z., Schiattarella, G. G., Bezprozvannaya, S., Raguimova, O. N., Cho, E. E., Vidal, A. H., Robia, S. L., Bassel-Duby, R., and Olson, E. N. (2018) The DWORF micropeptide enhances contractility and prevents heart failure in a mouse model of dilated cardiomyopathy. Elife 7

22. Singh, D. R., Dalton, M. P., Cho, E. E., Pribadi, M. P., Zak, T. J., Seflova, J., Makarewich, C. A., Olson, E. N., and Robia, S. L. (2019) Newly Discovered Micropeptide Regulators of SERCA Form Oligomers but Bind to the Pump as Monomers. J Mol Biol 431, 4429–4443

23. Schaaf, T. M., Li, A., Grant, B. D., Peterson, K., Yuen, S., Bawaskar, P., Kleinboehl, E., Li, J., Thomas, D. D., and Gillispie, G. D. (2018) Red-Shifted FRET Biosensors for High-Throughput Fluorescence Lifetime Screening. Biosensors (Basel) 8

24. Schaaf, T. M., Kleinboehl, E., Yuen, S. L., Roelike, L. N., Svensson, B., Thompson, A. R., Cornea, R. L., and Thomas, D. D. (2020) Live-Cell Cardiac-Specific High-Throughput Screening Platform for Drug-Like Molecules that Enhance Ca(2+) Transport. Cells 9

25. McCarthy, M. R., Savich, Y., Cornea, R. L., and Thomas, D. D. (2020) Resolved Structural States of Calmodulin in Regulation of Skeletal Muscle Calcium Release. Biophys J 118, 1090–1100

26. Li, J., James, Z. M., Dong, X., Karim, C. B., and Thomas, D. D. (2012) Structural and functional dynamics of an integral membrane protein complex modulated by lipid headgroup charge. J Mol Biol 418, 379–389

27. Muretta, J. M., Rohde, J. A., Johnsrud, D. O., Cornea, S., and Thomas, D. D. (2015) Direct real-time detection of the structural and biochemical events in the myosin power stroke. Proc Natl Acad Sci U S A 112, 14272–14277

